# Single-cell RNA-seq reveals TCR clonal expansion and a high frequency of transcriptionally distinct double-negative T cells in NOD mice

**DOI:** 10.1101/2023.07.21.550036

**Authors:** Md Zohorul Islam, Sam Zimmerman, Jon Weidanz, Jose Ordovas-Montanes, Michael Robben, Jacob M. Luber, Aleksandar D Kostic

## Abstract

T cells primarily drive the autoimmune destruction of pancreatic beta cells in Type 1 diabetes (T1D). However, the profound yet uncharacterized diversity of the T cell populations in vivo has hindered obtaining a clear picture of the T cell changes that occur longitudinally during T1D onset. This study aimed to identify T cell clonal expansion and distinct transcriptomic signatures associated with T1D progression in Non-Obese Diabetic (NOD) mice. Here we profiled the transcriptome and T cell receptor (TCR) repertoire of T cells at single-cell resolution from longitudinally collected peripheral blood and pancreatic islets of NOD mice using single-cell RNA sequencing technology. Surprisingly, we detected a considerable high frequency of islet-matching T cell clones in the peripheral circulation and blood-matching T cell clones in the islets. Our analysis showed that transcriptional signatures of the T cells are associated with the matching status of the T cells, suggesting potential future applications as a marker for early prediction of diabetes onset using peripheral T cells. In addition, we discovered a high frequency of transcriptionally distinct double negative (DN) T cells that might arise from naïve and effector backgrounds through the loss of CD4 or CD8 in a yet unknown biological pathway. This study provides a single-cell level transcriptome and TCR repertoire atlas of T cells in NOD mice and opens the door for more research into the causes of type 1 diabetes and inflammatory autoimmune disease using mouse models.

## Introduction

The onset of T1D results from the destruction of pancreatic β-cells by a multi-stage progression of autoimmune reactions. The islets remain free from lymphocytic infiltration in NOD mice until 3-4 weeks of age [1]. Gradually, lymphocytic infiltration starts with the sign of nondestructive insulitis, which is a hallmark of T1D pathogenesis [2]. At around 12 weeks, self-tolerance becomes broken due to the imbalance of regulatory T cells (Tregs) and effector CD8+ T cells resulting in targeted destruction of β-cells and onset of diabetes [3]. Insulitis in NOD mice involves many types of lymphocytes and myeloid cells [4,5]. However, the T1D pathogenesis observed in NOD mice is primarily driven by T cells [6–8]. It has been previously shown that CD4+ and CD8+ T cells are responsible for the autoimmune destruction of β-cells and diabetes development in these mouse models [9–11]. β-cell reactive T cells that target autoantigens like GAD, IGRP, and insulin antigen are commonly seen in T1D development of the NOD mice [12–16]. Other cell types such as B cells, dendritic cells, macrophages, and NK cells can also be found at the site of insulitis [17,18]. Once started, the insulitic lesion is continually replenished by new lymphocytes, mainly CD4+ and CD8+ T cells, from peripheral circulation [19].

Therefore, T cell recruitment to the islets is considered a continuing process. The appearance of autoreactive T cells in peripheral circulation during the progression of T1D is a critical marker for predicting the onset of diabetes. Previous studies demonstrated the predictability of T1D onset based on detecting autoreactive T cells in circulating peripheral blood [15,20,21]. T cell receptor (TCR) repertoire profiling is a powerful tool for identifying T cell clones and phenotypes directly linked to insulitis and β-cell destruction. TCR profiling can be performed by determining the sequence rearrangement of TCR αβ chains. Clonal expansion of T cells to pancreatic tissue and their linkage with the same circulatory clone in the blood can be detected by TCR profiling. However, most TCR profiling studies are limited to circulatory T cell clones [22–25] with few studies detecting clonal expansion of T cells in islets or other lymphoid organs, [26,27]. However, the majority of these studies were limited to relatively small sample sizes when compared to the theoretical biodiversity of TCR sequences. Other studies investigated the TCR clonal diversity in few select subpopulations of T cells within the pancreas and other lymphoid tissues in NOD mice, including CD4+ memory T cells [28,29] and T regs [30]. Marrero et al. [29] analyzed islet-infiltrating memory CD4+ T cells in prediabetic and recent onset diabetic NOD mice. They found many unique TCR clonotypes in islet-infiltrating CD4+ T cells and that TCR β repertoires were highly diverse at both stages of T1D development. While numerous studies reported a diverse TCR repertoire in peripheral blood and some other lymphoid organs in T1D patients, the diversity and clonal expansion of the T cell populations in pancreatic islets duringT1D onset is still unexplored. Recent technologies have enabled profiling the transcriptome at single-cell resolution, which has resulted in the growth of single-RNA sequencing studies of T1D tissues over the last few years [18,31–38]. Similar technologies enable the sequencing of TCR rearrangements at the level of individual clones, which can be collected concomitantly with RNA-sequencing data [39]. This study reports on the identification of T cell clonal expansion and distinct transcriptomic signatures that indicate progression towards T1D in the NOD mouse model and build on our predictive findings to identify T cell-intrinsic mechanisms that indicate disease progression.

## Results

### Single-cell RNA-seq identifies diverse T cell populations in peripheral blood and islets of NOD mice

We conducted paired single-cell RNA sequencing (scRNA-seq) and TCR repertoire sequencing on T cells collected from the peripheral blood and pancreatic islets of NOD mice (Figure 1a). We monitored female NOD mice from the age of 3 weeks to 40 weeks for the natural onset of diabetes (two consecutive blood glucose readings ≥250 mg/dl). We observed diabetic onset in 70% of NOD mice at 40 weeks of age without any significant difference in body weight between diabetic and non-diabetic groups (Figure 1b-1c). A total of six pooled-PMBC samples (collected a week before onset of diabetes) and their corresponding six pooled-islet samples (collected within one week of onset of diabetes) from diabetic mice were used for single-cell analysis (Supplementary file 1). Each pool consisted of three mice with diabetes onset at a similar time. In addition, four pooled PBMC samples from non-diabetic mice were collected at 34 weeks of age and included for comparison with PBMC of diabetic mice. The non-diabetic mice were monitored for diabetes onset until 40 weeks of age to be confirmed as free from hyperglycemia. There was no significant difference in T cell frequency in diabetic and non-diabetic mice blood (Figure 1d, Supplementary file 2). We performed a series of quality control steps during 10X library preparation (Supplementary Figure 1 and Supplementary Figure 2). As a result, we obtained 102,848 high-quality T cells from peripheral blood and islets (Supplementary File 2) that were included in the downstream analysis.

**Figure 1.**
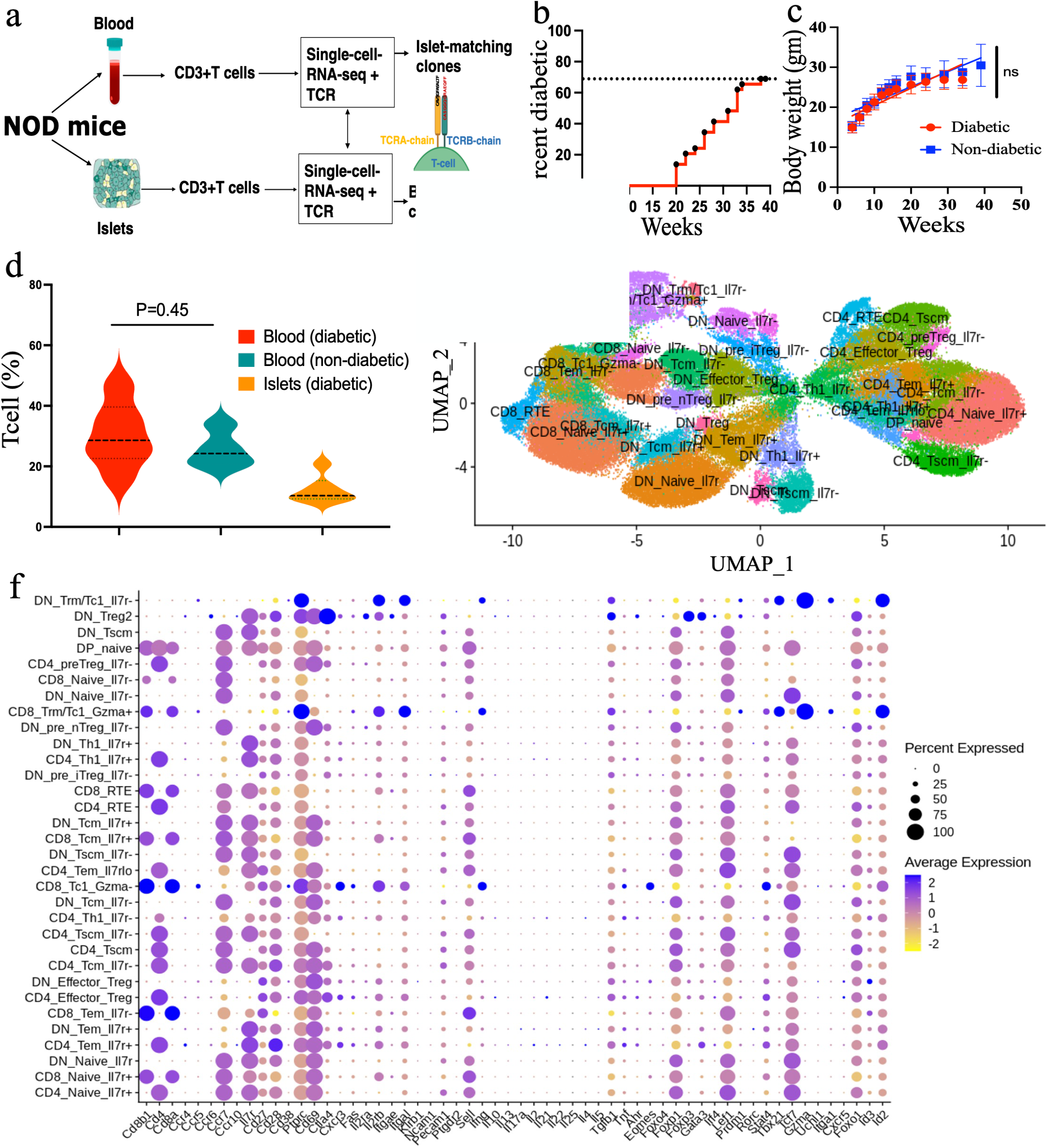
Single-cell RNA-seq analysis of peripheral blood and islet-infiltrating T cells from NOD mice. a. Experiment design for single-cell analysis of T cells and TCR clonality detection from peripheral blood and islets of NOD mice, including magnetic separation of CD3+ T cells, 5’-scRNA-seq, and V(D)J profiling, and computational approaches for TCR matching clone detection. b. Diabetes incidence and body weight growth in NOD mice in this study (c). Two consecutive blood glucose readings ≥250 mg/dl were considered for the onset of diabetes. The p values for the body weight difference between diabetic and non-diabetic mice indicate the non-significant difference between slopes (linear regression). d. Frequency of T cells among all cells in peripheral blood and islets. T cells were isolated by magnetic separation using monoclonal anti-mouse CD3ε antibody conjugated to Biotin. e. UMAP visualization of T cell populations in NOD mice. This UMAP represents T cells from the peripheral blood of diabetic mice (n = 46,741 cells) at the onset of diabetes, peripheral blood of non-diabetic NOD mice (n = 28,457 cells) at 34 weeks of age, and islet-infiltrating T cells (n = 27,650 cells) of diabetic mice at the onset of diabetes. f. Expression of canonical marker genes of T cells. Dot plots show the average and per cent expression of canonical marker genes across different T cell subpopulations. The cell clusters were annotated based on the expression of these marker genes.

We identified 32 T cell subpopulations (Figure 1e) in the peripheral blood and islets of NOD mice. T cell clusters were annotated based on the expression patterns of canonical marker genes (Figure 1f, Figure 2a-2d, Supplementary File 3, Supplementary Figure 3). Most islet-infiltrating T cells clustered separately compared with the T cells from peripheral blood (Figure 2e). We found a high rate (∼33%) of double negative (DN) T-cells (Figure 2f), lacking expression of either CD4 or CD8, among diabetic and non-diabetic mice in all tissue types, which is consistent with other reports [40]. This affected cell clusters, leading to the identification of clusters of CD4-CD8-Naïve, Tem, and Treg populations (Figure 2g). DN T cell populations displayed similar representation to CD4+ T cells in all tissue types, where CD4+ and DN Tregs were greater represented in islets than CD8+ Tregs (Figure 2g). We also observed specificity in diabetic mouse pancreatic islet samples towards CD8+ Tem and CD8+ Gzma-Tc1s (Figure 2e). We analyzed six paired islet pools from the same diabetic mouse cohort to identify the islet-infiltrating T cell landscape at the single-cell level. We found that islet-infiltrating T cells are dominated by diverse effector cell types (Figure 2g). The most frequent subtypes were CD8 Tem IL7r-, CD4 Th1 and CD4 effector Tregs (Figure 2g).

**Figure 2.**
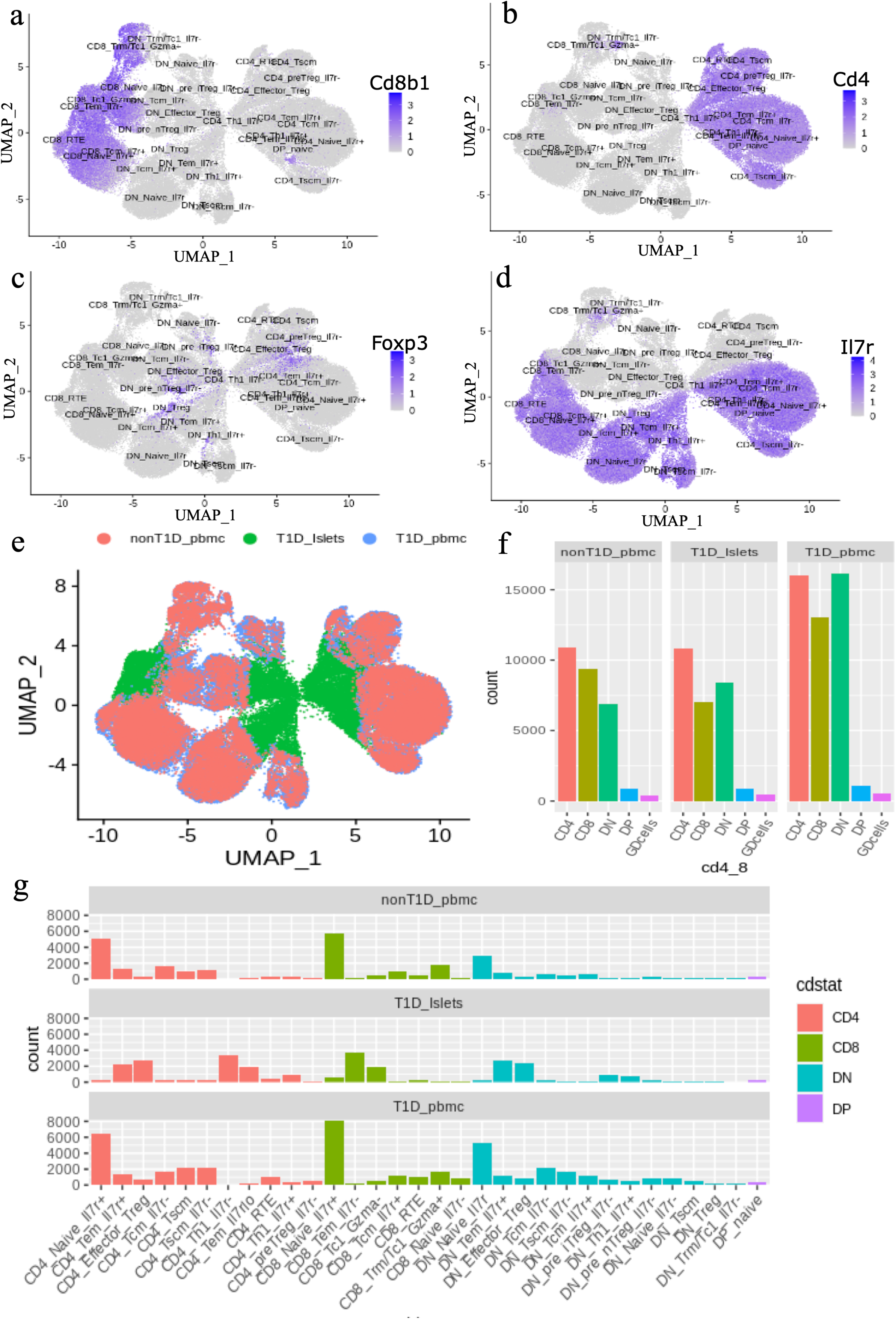
Frequency of T cell subpopulations in blood and islets. a-d. Feature plots visualizing the expression of major T cell markers CD4 and CD8, regulatory T cell marker Foxp3, and activation marker IL7r. e. UMAP visualization of T cell populations across three sample types. It shows a clear separation of T cell populations between islets and peripheral blood. f. Bar plot showing a high frequency of double-negative (CD4- and CD8-) T cells in all sample types. g. T cell subpopulation frequency in the peripheral blood of non-diabetic mice (top), islets of diabetic mice (middle), and peripheral blood of diabetic mice (bottom).

### Clonal expansion of islet-matching T cells can be detected in peripheral blood

We assessed clonal overlap and clonal expansion of T cells between blood and islets. First, we compared the T cells from paired blood and islets to identify an identical T cell clone in blood and islets from the same mouse pool. If a T cell in blood carries identical TCR α and β chains as another T cell in islets, it is defined as an “islet-matching” (IM) T cell clone in blood. By contrast, if a T cell in an islet carries identical TCR α and β chains with another T cell in the blood, it is defined as a “blood-matching” (BM) T cell clone in islets. Analyzing paired blood and islet samples from diabetic NOD mice, we were able to trace clones of islet-infiltrating T cells in the blood (Figure 3a). Only about 19% of the T cells sequenced had matching TCR sequences between blood and islet samples, but some populations were greater represented than others. For instance, we found that more CD8+ Gzma-Tc1s were blood matched than any other population and represented twice as much as CD8+Gzma+ Tc1s (Figure 3a and Figure 3b). We found that CD4+ and DN effector memory and effector Tregs were overrepresented among blood-matched T cells. CD4+ and DN Th1 cells were also found in both blood and islet samples.

**Figure 3.**
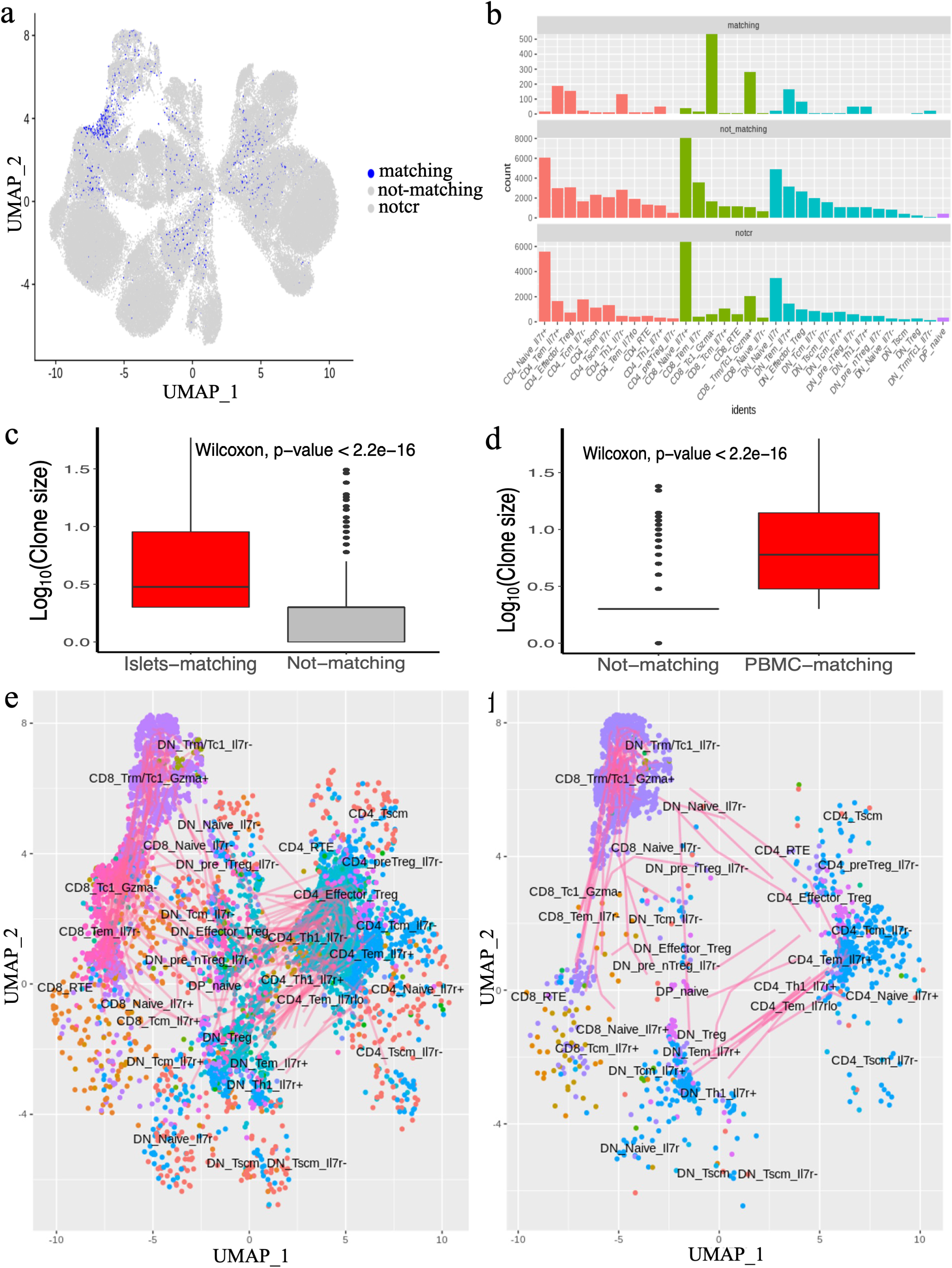
Clonal expansion of T cells in NOD mice. a. UMAP showing matching T cell clones in islets and peripheral blood. b. Frequency of different T cell populations based on the TCR matching status of T cell clones. The matching status of the T cell clones was defined if T cells in blood share identical TCR sequences (alpha/beta or both) with T cells in islets. c. Box plot quantifying clonal expansion of islet-matching T cell clones in blood. Boxes show the first, median, and third quartile, and the whiskers show 1.5× the interquartile range. Based on the Wilcoxon rank-sum test, p values show a statistically significant difference in clonal expansion between islet-matching and non-matching cells. d. Box plot quantifying clonal expansion of blood-matching T cell clones in islets. e. TCR clonal expansion of T cells (>4 cells) in peripheral blood and islets of NOD mice. f. TCR clonal expansion of T cells (>4 cells) in peripheral blood of non-diabetic NOD mice.

We then assessed the clonal expansion of T cells in the blood (Figure 3c). We found that the matching clones expanded significantly (P < 2.2X10^-16^) higher than the non-matching clones in the blood (Figure 3c). Next, we examined the clonal expansion of islet-infiltrating T cells with identical TCR in blood, termed BM T cell clones. BM T cells were present in almost all T cell subpopulations in islets (Figure 3b). Similar to IM clones in blood, the BM clones were found to be significantly (P < 2.2X10^-16^) highly expanded relative to non-matching cells in the islets (Figure 3d). Next, we investigated patterns of TCR clonal expansion in the peripheral blood and pancreatic islets of diabetic and non-diabetic mice. We observed a large amount of clonal expansion for diabetic Tc1 cells where cells localized to pancreatic islets experienced reduced expression of Granzyme A than circulating Tc1s (Figure 3e and Supplementary Figure 4). This was not seen to as great an extent in the non-diabetic mice (Figure 3f). We also found greater differentiation between clonal CD4+ and DN Tems and Tregs in diabetic mice.

### The matching clones of T cells show enriched interferon-gamma response pathways compared with non-matching clones

Although the matching clones in blood and islets carry similar TCR sequences, their transcriptional signatures differ. We identified 160 differentially expressed (DE) genes (Log fold-change of 1 or more) between matching cells in the blood and islets (Figure 4a, Supplementary File 4). DE analysis shows the BM cells in islets expressed elevated levels of effector and migration-related genes such as *Ccl3*, whereas the IM cells in blood show higher expression of cytotoxicity-related genes such as *Gzma* (Figure 4a). Gene set enrichment analysis (GSEA) showed that the matching cells in islets were enriched for interferon-alpha and interferon-gamma response pathways (Figure 4d). We then analyzed the DE genes between IM clones and non-matching clones in blood and found 40 genes upregulated and 13 genes downregulated (Log fold-change of 1 or more). The IM cells in the blood showed elevated effector and cytotoxicity markers in comparison to non-matching cells such as *Ccl5*, *Gzmk*, and *Nkg7* (Figure 4b, Supplementary file 5). The IL-2 STAT5 signaling pathways were more significantly enriched in the matching cells than in non-matching cells in the blood (Figure 4e). In addition, the interferon-gamma response pathway was elevated in the matching cells. Similar to blood, the matching cells in the islets showed elevated expression levels of effector and cytotoxicity markers such as *Ccl5*, *Ccl4*, *Gzmk*, and *Nkg7* (Figure 4c, Supplementary File 6). The matching clones of T cells showed enriched IL-2 STAT5 signaling and interferon-gamma response pathways compared with non-matching clones (Figure 4f, Figure 4g). In addition, we observed significantly higher expression of *Nkg7*, Gzmk, and *Infg* genes in the matching cells compared with non-matching cells in the islets (Figure 4h). Furthermore, the non-matching cells in the islets showed significantly high *InfgR2* (interferon-gamma receptor) gene (Figure 4h).

**Figure 4.**
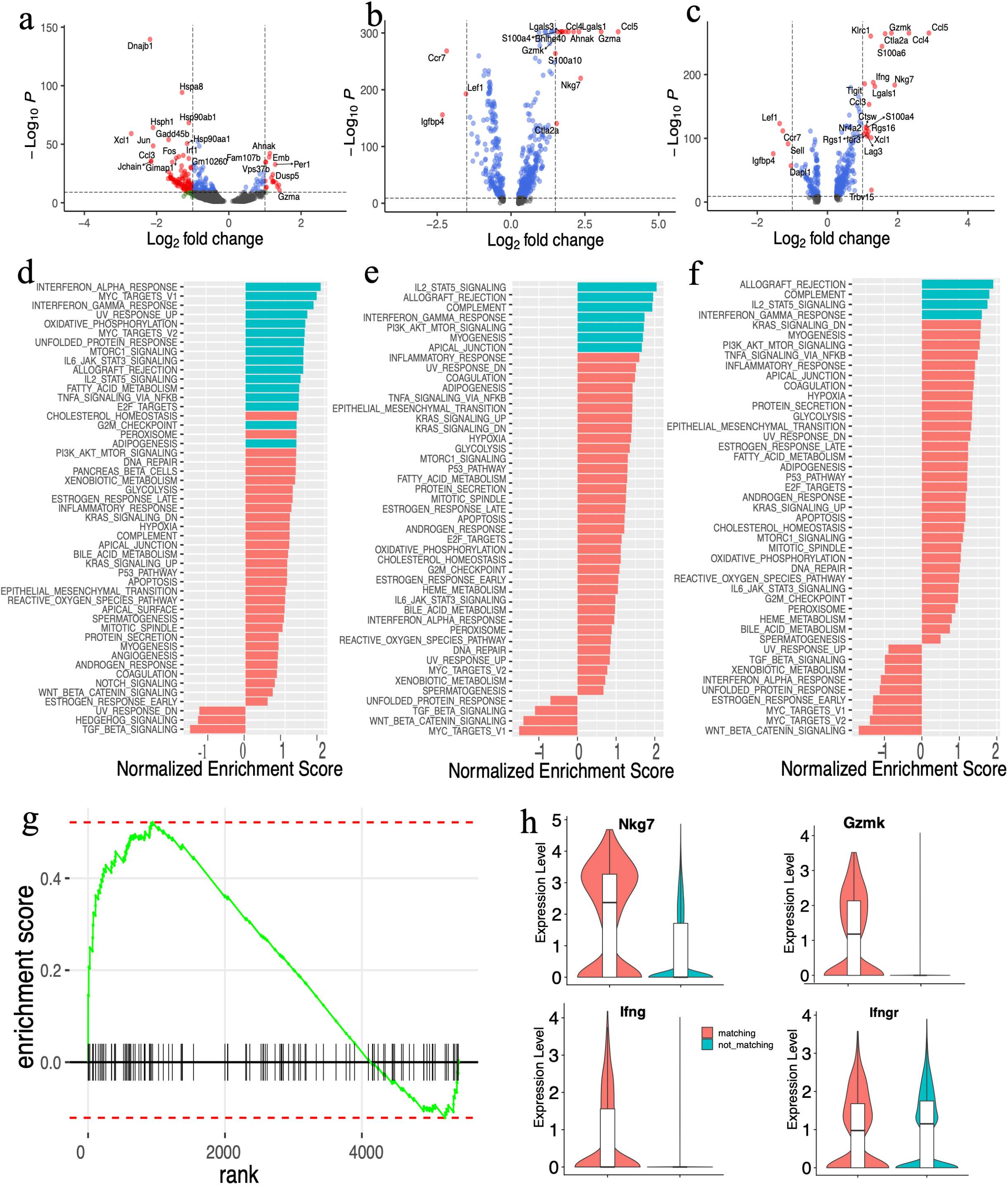
Transcriptional heterogeneity among matching T cell clones in blood and islets. Differential expression analysis of genes between the matching clones in islets and blood (a), islet-matching and non-matching clones in blood (b), and non-matching and blood-matching clones in islets (c). d. Gene Set Enrichment Analysis (GSEA) shows enriched pathways in matching clones in islets compared with the matching clones in blood. Significantly enriched pathways (padj <0.05) are highlighted with green colors. HALLMARK pathway set from Msigdb database was used for the GSEA. e. GSEA shows enriched pathways in islet-matching clones in blood compared with non-matching clones. f. GSEA shows enriched pathways in blood-matching clones in islets compared with non-matching clones. g. Enrichment plot showing the enrichment score of genes involved in the enrichment of interferon-gamma response pathway in matching clones compared with non-matching clones in islets. h. Significantly higher expression of cellular cytotoxic accelerating gene Nkg7 and Gzmk, and proinflammatory mediator Inf-g in matching cells compared with non-matching cells. It also shows increased expression of the interferon-gamma receptor (InfgR) gene by the non-matching cells.

### Transcriptome profiles of T cell clones are associated with the matching status

We observed that the matching cells are transcriptionally distinct from non-matching cells. We wanted to test whether a machine learning classifier could be trained to predict if a given T cell from blood is IM or non-matching based on the transcriptional signature. We predicted the matching and non-matching status based on the expression of genes in the CD8 cell compartment in T1D blood where we observed most of the matching cells. We demonstrated that a regularized logistic regression classifier achieved moderately high sensitivity and specificity (Figure 5a and Figure 5b, Overall area under the curve [AUC] = 0.89 and overall AUPRC = 0.67). Our analysis showed that transcriptional signatures of the T cells are associated with the matching status of the T cells, suggesting potential future applications as a marker for early prediction of diabetes onset using peripheral T cells. We then identified which genes are most likely to be associated with the matching status of the cells. Finally, we identified a list of genes that were found to be associated with the matching status of the cells (Figure 5c).

**Figure 5.**
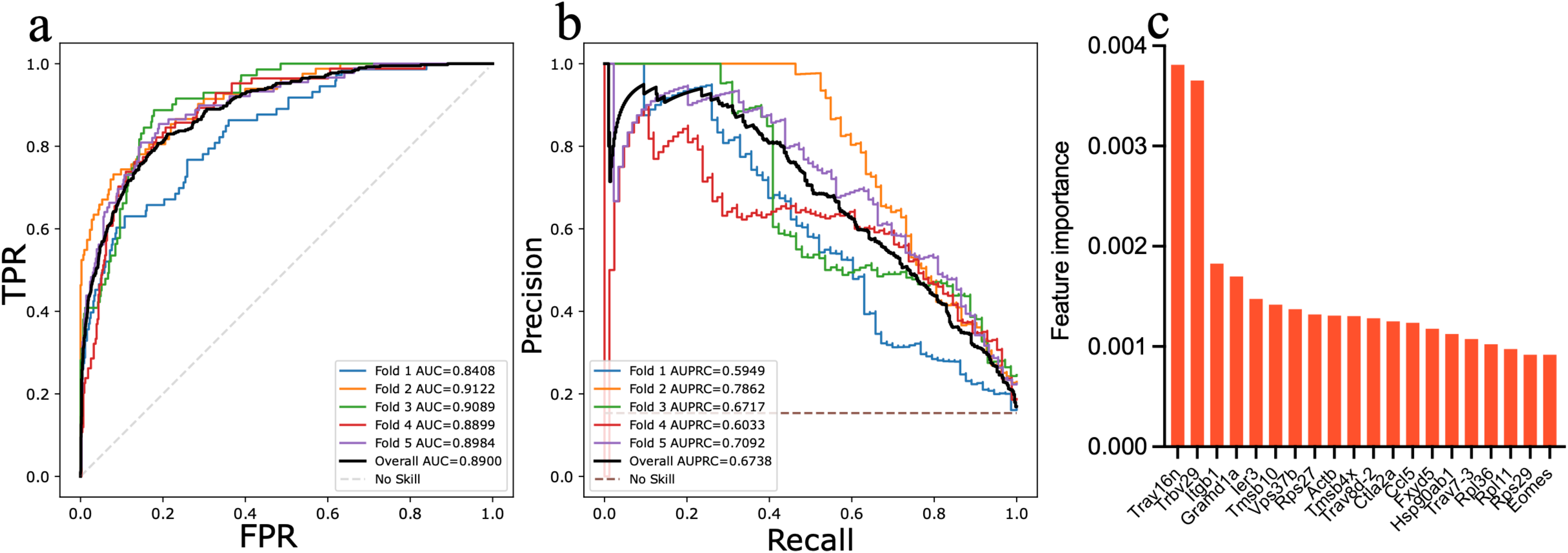
The transcriptional signatures of T cells can predict the matching status of the T cell clone. a-b. Receiver operating characteristic curve and recall-precision plot with five-fold cross-validation showing the classification of cells as IM or non-matching in the CD8 effector memory cells in blood based on logistic regression classifier. c. A bar chart with the top 20 genes was found to be the most significant predictor of the matching status of the cells. The Y-axis shows feature importance.

### Single-cell trajectory analysis reveals the enrichment of T cell cluster-specific gene module

Our results showed that the transcriptional profile of T cells’ matching clones in the islets is distinct from non-matching cells. We observed that most of the matching clones were found in cell compartments, such as the CD8 Tc1 GZMA-cluster. We wanted to detect differentiation patterns within our member T cell populations, so we used the monocle3 pipeline to derive psuedotime calculations for each cell (Figure 6a and Figure 6b). Psuedotime analysis shows a logical progression from Naïve T cell populations to T effector memory and effector Tregs. We found that DN Tregs have unique naïve populations that follow similar lymphocyte proliferation patterns. An interesting result is that Naive CD8+ IL7R-T cells can differentiate into DN effector cells and Tregs through a previously unknown pathway. We also observed possible state transitions from DN T cells to T effector populations, yet this pathway is also uncharacterized.

**Figure 6.**
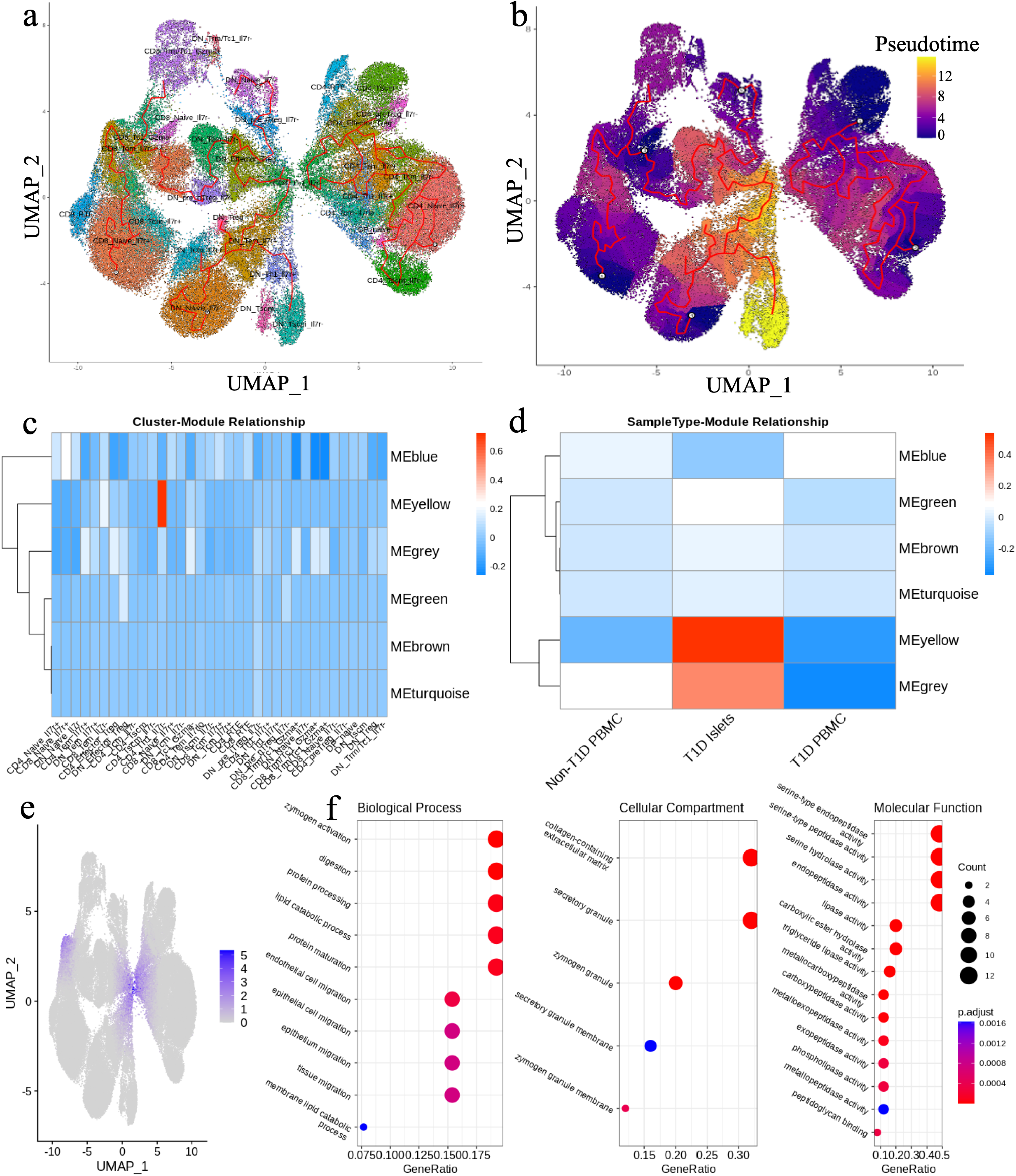
T cell trajectory and weighted gene co-expression network analysis (WCGNA) of islet-infiltrating and peripheral blood-derived T cells of NOD mice. Transcriptional trajectory **(a)** and pseudo temporal axis **(b)** of T cell across different subpopulations. Employing WCGNA, we identified a specific gene module among T cell populations **(c)** and sample types **(d)**. **e**. Feature plots showing the expression of yellow modules. The scale bar indicates the expression level. **f.** Gene Ontology (GO) enrichment analysis shows the biological functions of the genes in the yellow modules. The color scale represents Benjamini-Hochberg adjusted p-values, and the size of the circles means gene count.

Progression of a cell state can also be identified through modules of genes that express similarly to each other. A module specific to granzyme regulation (MEYellow) is found enriched and expressed at higher rates in CD8+ Gzma-Tc1 cells and DN/CD4+ Th1 cells but not in CD8+ Gzma+ Tc1 cells (Figure 6c). Interestingly, the MEYellow module was mostly enriched in islets infiltrating cells (Figure 6f). Further, we performed Gene Ontology (GO) enrichment analysis to picture the biological functions of the genes in the yellow modules. We found that zymogen activation pathways related genes were highly enriched in this module (Figure 6f). On the other hand, the DN Tregs show higher expression the MEgreen module (Figure 6c, Supplementary Figure 5b). We also observed expression of a module associated with MHC processing and binding expressed greater only in DN Treg cells and their precursors (Supplementary Figure 5b).

## Discussion

The discovery of biomarkers for early detection of T1D is a pressing need in the medical community. It will be transformative for diagnosing and treating the growing population affected by T1D [41]. Combined single-cell transcriptome and TCR repertoire analysis provide the opportunity to dissect the functional diversity of clonally expanded T cells. This study identified clonal expansion and transcriptional heterogeneity of T cells in peripheral circulation and pancreatic islets in NOD mice using paired scRNA-seq and TCR profiling. Clustering single-cell RNA-seq data to identify previously known or novel cell types has been shown to be challenging to perform in a way that reflects the actual biological state of cells due to inaccurate algorithms and a lack of a priori knowledge of cell-specific gene expression [42]. We chose to cluster cells in such a way that they maximized biological diversity between subsets of T cells, applying k-Nearest Neighbors (KNN) based clustering algorithms to weighted PCA that put more importance on the variance of immunologically relevant marker genes. We acknowledge that the resulting cell clusters may not be truly representative of the underlying biology, but we have developed several metrics to demonstrate to us that they are robust enough to analyze the patterns of expression across different types of T cells.

We identified a high-resolution T cell landscape in the peripheral blood and islets of NOD mice. The overall frequency of Th1 type effector cells was much higher in the islets than in the paired blood samples of diabetic mice, indicating the transcriptional transition of T cells upon entering islets from blood. This vast difference in T cell phenotypes between blood and islets raised the question of whether the influx of T cells into the inflammatory pancreatic lesion is a continuous process involving freshly recruited cells, as observed by an earlier study [19]. If the T cell recruitment to this site is a regular influx from the circulation, we could see overlapping cell phenotypes between blood and islets. The T cell populations in the islets sampled for this study display functional changes that suggest they differentiate and multiply locally after they infiltrate the pancreatic islets. For instance, the islets of diabetic mice show a substantial, large size Th1-like cell cluster that was not detected in the blood. This finding might indicate that some of the T cell types in the blood of diabetic mice undergo the transition to Th1-like cells upon entering the insulitis site.

Another interesting finding of this study is the presence of very high frequency transcriptionally distinct DN T cells in NOD mice. Although 1-5% of the T cells in rodents and humans are usually found as double negative [43], we detected a high frequency (33%) of double negative (DN) T cells both in islets and in peripheral blood of diabetic and non-diabetic NOD mice. This result was unexpected and yet several studies reported increased DN T cell representation in NOD mice [40,44]. Liu et al. [44] even showed that DN Tregs can rescue the non-diabetic phenotypes in diabetes-induced SCID mice that were transfected with NOD CD4+ Tregs. Within our sequenced population, we found that while DN T cells are transcriptionally distinct from CD4+ and CD8+ T cells, they show parallel cell types to those found in traditional T cells. The most surprising cell type we identified in our sample were CD4-CD8-Naïve T cells, which are theoretically not even able to be generated as a product of TCR processing in T lymphocyte development [45]. Because these T cells are present in both diabetic and non-diabetic mice, we believe that increased levels of DN T cells are a feature of the NOD genetic background that may lead to diabetes and autoimmunity. Based on psuedotime analysis, we also believe that these DN T cells arise from both naïve and effector backgrounds through the loss of CD8 or CD4 in a yet unknown biological pathway.

Surprisingly, we detected a considerable high frequency of IM T cell clones in the peripheral circulation and BM T cell clones in the islets. The matching clones in both tissues were found to be significantly expanded compared with non-matching clones, which raised questions about where this expansion occurs. The autoreactive T cell may become clonally expanded upon interacting with antigens presented by antigen-presenting cells in the pancreatic lymph node and enter the islets. However, this observation raised several questions about why there is a substantial amount of clonally expanded matching cells in the blood. Do these cells come back from the immunogenic islets to the blood, or does the expansion happen in other lymphoid tissues and circulate systematically? Differential gene expression and GSEA identified functional connections between the matching and non-matching cells in the islets. We assume the matching clones are the first invader to the islets. Once they are accumulated, the early-invading cells may attract other non-specific cells by secreting cytokines and chemokines signals and accelerating tissue destruction. It is supported by the increased expression of cytotoxic effector-like and chemokine signaling molecules by the matching cells. For instance, the BM cells in the islets express a significantly higher level of *Nkg7* than non-matching cells. Studies show that *Nkg7* accelerates cellular cytotoxicity and the production of inflammatory factors [46]. We found that the matching cells in islets express significantly high cytotoxic and inflammatory factors like *Gzmk*, *Ccl5* and *Infg*. The GSEA also shows that the interferon-gamma response pathways were significantly enriched in the matching cells compared with non-matching cells in islets. Further, the non-matching cells expressed a significantly high interferon-gamma receptor 2 (*InfgR2*) level from the matching cells, indicating their response to *Infg* signals. Our results highlight Nkg7 as a potential target of therapeutics to prevent the inflammatory response that leads to insulitis and T1D by reducing accumulation of autoreactive T cells in the pancreatic islets.

Applying a linear regression model to the data, we showed that the transcriptional profile of T cell clone could be used as a predictor of a cell’s matching status. Furthermore, we identified a set of genes that explains the matching and non-matching association. However, future research is required to determine how early this matching clone of T cells can be detected in the peripheral circulation and whether a similar gene set can explain the matching status.

A major finding in our results is that CD8+ Tc1 cells are downregulated for Granzyme A expression in the pancreatic islets. This suggests that something in the pancreatic islet is regulating Tc1 function to prevent cytotoxic activity related to diabetic progression. A recent study in Granzyme A deficient NOD mutants showed similar tolerance and systemic autoimmunity loss that enhances diabetes progression [47]. This was shown to be Ifng dependent, but interestingly we found that CD4+ Th1 cells native to the NOD pancreatic islets did not express Ifng (Figure 1f). We have no explanation currently for what regulates Granzyme A expression within CD8+ Tc1 cells nor what causes differentiation of CD8+ and CD4+ T cells to DN T cells or what role these things play in the development of T1D. In addition, we found a substantial large subtype of Th1 cells in the islets that were not detected in such a high proportion in blood. Despite the low abundance of Th1 cells in the blood, their abundance in the islets indicates that these cells might be continuously recruited or converted from other cell types in the presence of cytokines or chemokines signals [48]. A Previous study has shown that Th17 cells promote islet inflammation by conversion into Th1 cells [49] and IL-17 deficient NOD mice show delayed onset of diabetes [50]. Therefore, we wanted to know whether we could see any difference in the level of Il17a, a major cytokine secreted by Th17 cells, between diabetic and non-diabetic mice. We analyzed blood plasma samples at three weeks, 10 weeks, and at the terminal time point and found an elevated trend of IL17a in the blood of diabetic mice than in non-diabetic mice (Supplementary Figure 6a). However, we did not find any particular cell cluster showing distinct expression of IL17a (Supplementary Figure 6b). It is possibly due to the reason that we determined IL17a abundance in blood plasma through ELIZA whereas the single cell data shows the expression of the mRNA just through the enriched T cells. It could be a non-T cell source of IL17a or perhaps we just didn’t observe the expression because the expression stopped by the time, we harvested the cells.

While our study shares some similarities of study design with a recently published paper [51], it contributes valuable understanding to the T cell landscape in T1D. Both studies identified phenotypic and transcriptional differences in CD4+ and CD8+ T cell compartments in spontaneous onset diabetes in NOD mice. However, our research took a different approach by observing the spontaneous onset of diabetes in NOD mice until 40 weeks of age to confirm non-diabetic phenotype, allowing us to compare the T cell transcriptional heterogeneity between diabetic and non-diabetic mice, thus mimicking the natural biology. In contrast, the Collier et al. 2023 study [51] defined spontaneous diabetes onset at 20 weeks and compared it with an anti-PD1 treated accelerated onset diabetic mice cohort. Moreover, our study focused on the clonal expansion and transcriptional heterogeneity of T cells in the peripheral circulation and pancreatic islets of NOD mice. We discovered the presence of DN T cells and their potential role in T1D. On the other hand, the Collier et al. study emphasized the comparison between spontaneous and anti-PD-1-induced T1D and highlighted the importance of monitoring autoimmune diseases in peripheral tissues, such as blood. Furthermore, our research demonstrated the functional differences between matching and non-matching T cell clones in the islets and peripheral circulation, providing the dynamic behavior of T cell populations in T1D. While both studies contribute valuable understanding into the functional diversity of T cells, our comprehensive analysis offers a more detailed understanding of clonal expansion, transcriptional heterogeneity and phenotypic transitions with regard to natural onset diabetes in NOD mice.

## Conclusion

We characterized and identified a single-cell level T cell map in the peripheral blood and islets of NOD mice during the progression of Type 1 diabetes. We were able to locate islet-matching T cell clones in the blood and their expansion both in blood and islets. This study discovered a high frequency of transcriptionally distinct DN T cells that might arise from naïve and effector backgrounds through the loss of CD8 or CD4. We are sure that these discoveries have opened the door for more research into the causes of Type 1 diabetes and inflammatory autoimmune disease using mouse models.

## Materials and methods

### Experiment mice model

We purchased three-week-old female NOD/ShiLtJ mice from The Jackson Laboratory (stock number 001976) and were monitored for natural diabetes onset until 40 weeks of age. The mice experiment was carried out in a pathogen-free facility at Joslin Diabetes Center (JDC) under standard housing, feeding, and husbandry. The mice experiment protocol was reviewed and approved by JDC’s Institutional Animal Care and Use Committee (IACUC) (Protocol #2016-05). In addition, ARRIVE guidelines were strictly followed while conducting the mice experiment.

### Blood glucose measurement and diabetes diagnosis

Tail blood was used to measure glucose concentration. Every two weeks, blood glucose was measured by Infinity blood glucose test strips (GTIN/DI#885502-002000) and an Infinity meter. Mice showing two consecutive glucose readings of ≥250 mg/dl were considered diabetic.

### Blood lymphocyte collections and cryopreservation

Peripheral blood was collected from mice every two weeks. Approximately 200 ul of blood from each mouse was collected from the tail using a heparin-coated Microvette tube to prevent blood clots (Sarstedt catalog#16.443.100). The blood was gently homogenized inside the heparin blood collection tube and diluted the blood by adding 1 mL of RPMI+10% fetal bovine serum (RF10). The RF10 media was added to dilute the effect of anticoagulants. The diluted blood was then slowly overlayed at room temperature on top of an equal volume of Histopaque 1083 (Sigma-Aldrich, Missouri, USA catalog#1083-1) at the bottom of a 5mL polypropylene tube maintaining a sharp interface between the blood and Histopaque. Next, the blood-Histopaque mixture was centrifuged at 400xg for exactly 30 minutes at room temperature (RT) with slow acceleration and without a break. Centrifugation was performed at RT because lower temperatures, such as 4 °C, may result in cell clumping or poor recovery. After centrifugation, a 500 ul top plasma layer was carefully transferred to an Eppendorf tube without disturbing the middle white buffy-coat layer.

The plasma was preserved at -20 C for further cytokine measurements. The white buffy coat layer at the Histopaque-media interface (red blood cells go down to the very bottom of the tube) was transferred to a clean 5 mL polypropylene tube and added 2 mL RF10 media for washing. The cell suspension was mixed by gentle inversion of the tube several times and followed by centrifugation at 300xg for 10 minutes at 4 °C with brake and acceleration. The washing step was repeated three times. Total cell count and viability were performed by staining with 0.4% Trypan blue and counted by the Countess automated cell counter. After counting, the cell pellet was resuspended in 1 mL of Cryostor CS10 (Stemcell Technologies Catalog #07930) and transferred to a cryotube. The PBMC containing cryotubes were placed in a polycarbonate container (Nalgene® Mr. Frosty) for controlled freezing and kept at -80 °C for 48 hours, followed by long-term preservation in liquid nitrogen until further analysis.

### IL17a ELISA

Cryopreserved blood plasma was thawed at room temperature and used for IL17a detection by a commercial mouse IL-17a ELISA Kit (Sigma Aldrich catalog #RAB0263-1KT). The test was performed according to manufacturer guidelines. The concentration of IL17a was determined from a standard curve generated by the known concentration of IL17a positive control.

### Single-cell collection from pancreatic islets and cryopreservation

We sacrificed the mice at the onset of diabetes and harvested the pancreas by collagenase perfusion. Mice were anesthetized with a standard dose of ketamine/xylazine before initiating the surgery. By making a V-cut from the lower abdomen, the pancreas was opened as much as possible. The pancreatic duct at its duodenal insertion was clamped off with a micro clamp, and 3 mL of freshly prepared collagenase enzyme (Vitacyte, CIzyme catalog #005-1030) was injected through the bile duct at the proximal end to perfuse the enzyme to the pancreas. After the collagenase infiltration, the pancreas was carefully collected in a 50 mL conical tube and incubated at 37 °C for 17.5 minutes. The incubation with collagenase was terminated by adding 25 mL cold RPMI+10% calf serum (RN10) media. The pancreas tissue was hand-shaken vigorously for 5-10 seconds and centrifuged at 500xg for 1.5 minutes (RT). This washing step was repeated two more times. The tissue suspension was resuspended in 25 ml of RN10 media and filtered through a 425 um diameter wire mesh (Cole Parmer catalog #SI59987-16) to remove the remaining undigested tissue, fat, and lymph nodes. The filtrate was centrifuged at 500xg for 1.5 min at RT, and the tissue pellet was resuspended in 10 mL lymphocyte separation media (LSM) (Corning, cat #25072-CV) by gentle vortex until the suspension was homogeneous. The tissue-LSM suspension was overlaid with 5 mL RN10 media to make a density gradient mixture, being careful to maintain the sharp interface between the LSM and the media. The density gradient mixture was centrifuged at 2500xg for 17 minutes at 4 °C with slow acceleration and no braking. After centrifugation, the islets layer at the interface (10mL) was collected in 15 mL RN10 media and centrifuged at 500xg for 2 min at 4 °C. Subsequently, islets were washed two times with 25mL RN10 media at 500xg for 2 min at 4 °C. Non-islet tissues were removed from the islets suspension by microscopic examination. These purified islets were subjected to non-enzymatic single-cell dissociation. First, The islets suspension was centrifuged at 500xg for 1.5 min at 4 °C. Then, the pellets of islets were resuspended in 2 mL warm non-enzymatic cell dissociation solution (Sigma-Aldrich, catalog##C5789-100ML) and incubated at 37 °C for 3 minutes. Islets were dispersed into single cells with frequent pipetting using wide-bore pipette tips. The cell suspension was centrifuged at 500xg for 5 minutes at 4 °C and resuspended in 5 mL of cold RPMI+10% FBS (RF10) media. The cell suspension was filtered through a 70μm cell strainer and counted before cryopreservation as mentioned in the previous section (see the cryopreservation method for PBMC).

### T cell preparation for single-cell RNA-sequencing

PBMC and islets single-cell suspension preserved in liquid nitrogen were revived before the magnetic separation of CD3+ T cells. The cryopreserved samples were processed according to the guidelines of 10x genomics (Document CG00039 Rev D) with slight modification. Briefly, the cryovials containing samples were immediately thawed in a water bath at 37 °C for 2-3 min and the cells were slowly transferred to a 50 mL Falcon tube using a wide-bore pipette tip.

Sequentially, the cells were gradually diluted to 32 mL RF10 media by incremental 1:1 volume additions of media for a total of 5 times. The cell suspension was then centrifuged at 300xg for 5 minutes at RT and resuspended in 10 mL RF10 media. Finally, the cells were washed twice in 10 mL RF10 media and centrifuged at 300xg for 5 minutes at RT to remove the remnants of cryoprotectants. Before proceeding to CD3+ cell enrichment, dead cells and debris were removed using Miltenyi MACS Dead Cell Removal Kit (Miltenyi catalog #130-090-101), following manufacturer guidelines. After dead cell removal, the CD3+ T cells were isolated using the Miltenyi MACS CD3 microbead kit (Miltenyi catalog#130-094-973) following the manufacturer and 10x genomics recommended guidelines (10x Protocol CG000123 Rev B). The cell viability and count were performed before and after CD3+ T cell enrichment. The magnetically separated CD3+ T cells were placed on ice and immediately used for single-cell cDNA library preparation.

### Single-cell RNA and immune repertoire sequencing library preparation

Single-cell 5’ gene expression (GEX) and V(D)J sequencing libraries of T cells were generated using the Chromium Next GEM Single Cell 5’ Dual Index Reagent Kits v2 (10x Genomics, Pleasanton, CA, USA) following the manufacturer guidelines (10X protocol CG000331 Rev A). All necessary primers and reagents were supplied within the 10x genomics reagent kits. The key steps involved in library preparation are briefly summarized below.

### GEM generation, barcoding, and cDNA synthesis

Single-cell suspensions of CD3+ T cells were diluted with the appropriate volume of nuclease-free water to achieve a targeted recovery of 10,000 cells per sample. The diluted single-cell suspension was mixed with Master Mix and loaded onto the 10x microfluidic chip Chromium Next GEM Chip K with combined barcoded Single Cell VDJ 5’ Gel Beads and Partitioning Oil. Cells were partitioned into nanoliter-scale Gel Beads-in-emulsion (GEMs) in Chromium Single Cell Controller using the Single Cell 5’ v2 protocol. Successful GEM generation was ensured by visual inspection of homogeneity of volumes and colors of GEM suspension across multiple samples. Next, GEM-RT incubation was performed to produce 10x Barcoded, full-length cDNA from poly-adenylated mRNA. GEMs was incubated on Bio-Rad C1000 Touch thermal cycler with two steps of incubation at 53 °C for 45 min in step 1 and 85 °C for 5 min in step 2.

Dynabeads MyOne SILANE magnetic beads were used to clean up the 10x Barcoded first-strand cDNA after the GEM-RT incubation step. Finally, 10x Barcoded, cDNA was amplified by PCR with primers against common 5’ and 3’ ends added during GEM-RT incubation. The cDNA amplifying PCR was performed with initial one step denaturation at 98 °C for 45Ls, followed by 13 cycles of denaturation (98 °C for 20Ls), annealing (67 °C for 30Ls) and extension (72 °C for 1 min), one step final extension at 72 °C for 1 min, and final holding at 4 °C. The amplified cDNA was cleaned-up using SPRIselect paramagnetic beads (Beckman Coulter, Brea, CA, USA) and freshly prepared 80% ethanol. Quality control and quantification of cDNA were performed using 2100 expert High Sensitivity DNA Assay (Agilent Technologies, Santa Clara, CA, USA). This quality-controlled cDNA was used for single-cell 5’ gene expression and VDJ sequencing libraries.

### Construction of single-cell 5’ gene expression library

The concentration of the amplified cDNA was adjusted to 50 ng per 20 ul volume in each sample to be used for 5’GEX library preparation. First, the cDNA amplicon size was optimized by enzymatic fragmentation, end repair, A-tailing, and double-sided size selection. The cDNA and 10x fragmentation enzyme mix were incubated in a pre-cooled thermal cycler at 32 °C for 5 min for fragmentation, followed by end-repair and A-tailing reaction at 65 °C for 30 min with a final hold at 4 °C. The fragmented cDNA was purified by double-sided size selection using SPRIselect beads. Second, the adapter oligos (P5 and P7) were ligated with the cDNA fragments and cleaned up. The adapter ligation reaction was performed at 20 °C for 15 min with a final hold at 4 °C. Post ligation cleanup was performed using SPRIselect beads. Finally, each sample was dual indexed with i5 and i7 sample index (PN-3000431 Dual Index Plate TT Set A, 10x Genomics, Pleasanton, CA, USA) using index PCR followed by cleanup and library QC. The indexed PCR was performed with initial one step denaturation at 98 °C for 45Ls followed by 14 cycles of denaturation (98 °C for 20Ls), annealing (54 °C for 30Ls) and extension (72 °C for 20 s), one step final extension at 72 °C for 20 s, and final holding at 4 °C. The sample indexed libraries were cleaned up using SPRIselect beads. 5’GEX Library quality control and quantification were performed using 2100 expert High Sensitivity DNA Assay.

### TCR enrichment and construction of V(D)J library

The cDNA was used to amplify full-length V(D)J segments with 10x Barcode via PCR amplification with primers specific to the TCR constant region. Using 2 ul cDNA of each sample, the V(D)J enrichment was performed by two rounds of PCR amplification. The first PCR amplification was performed using mouse T Cell Mix 1 v2 reagent (PN 2000256, 10x Genomics) that involved one step denaturation at 98 °C for 45Ls followed by 12 cycles of denaturation (98 °C for 20Ls), annealing (62 °C for 30Ls) and extension (72 °C for 1 m), one step final extension at 72 °C for 1 m, and final holding at 4 °C. The first V(D)J amplification PCR products were purified by double-sided size selection using SPRIselect beads before conducting the second round of PCR. The second PCR amplification was performed using mouse T Cell Mix 2 v2 reagent (PN 2000257, 10x Genomics) that involved one step denaturation at 98 °C for 45Ls followed by 10 cycles of denaturation (98 °C for 20Ls), annealing (62 °C for 30Ls) and extension (72 °C for 1 m), one step final extension at 72 °C for 1 m, and final holding at 4 °C. The V(D)J PCR amplicons were purified using SPRIselect beads before V(D)J library construction. Quality control and quantification of V(D)J amplicons were performed using 2100 expert High Sensitivity DNA Assay. The concentration of the V(D)J amplicons was adjusted to 50 ng per 20 ul volume in each sample to be used for V(D)J library preparation. First, the V(D)J amplicon size was optimized by enzymatic fragmentation, end repair, A-tailing, and double-sided size selection. The V(D)J amplicons and 10x fragmentation enzyme mix were incubated in a pre-cooled thermal cycler at 32 °C for 2Lmin for fragmentation, followed by end-repair and A-tailing reaction at 65 °C for 30 min with a final hold at 4 °C. SPRIselect beads purified the fragmented V(D)J amplicons. Second, the adapter oligos (P5 and P7) were ligated with the V(D)J amplicon fragments and cleaned up. The adapter ligation reaction was performed at 20 °C for 15 min with a final hold at 4 °C. Post ligation cleanup was performed using SPRIselect beads. Finally, each sample was dual indexed with i5 and i7 sample index (PN-3000431 Dual Index Plate TT Set A, 10x Genomics, Pleasanton, CA, USA) using index PCR followed by cleanup and library QC. The indexed PCR was performed with initial one step denaturation at 98 °C for 45Ls followed by 8 cycles of denaturation (98 °C for 20Ls), annealing (54 °C for 30Ls) and extension (72 °C for 20 s), one step final extension at 72 °C for 20 s, and final holding at 4 °C. The sample indexed libraries were cleaned up using SPRIselect beads. V(D)J Library quality control and quantification were performed using 2100 expert High Sensitivity DNA Assay.

### Sequencing of 5’GEX and V(D)J libraries

Gene expression and V(D)J libraries were sequenced on Illumina NovaSeq6000 sequencer by GENEWIZ (GENEWIZ, LLC, NJ, USA). Each of the 5’GEX and V(D)J libraries were prepared with unique dual indexes, which allows multiplexing of all libraries for sequencing. Libraries were sequenced according to the 10X Genomics configuration. A minimum of 20,000 read pairs per cell were sequenced for 5’GEX libraries, and a minimum of 5,000 read pairs were sequenced for V(D)J libraries.

### Sequence demultiplexing and processing

Raw sequencing reads were processed using Cell Ranger version 6.0.2 to produce gene expression count matrices and TCR clonotype summary. Using the raw sequencing FASTQ files, we first run the *cellranger multi* pipelines for simultaneous processing and analysis of V(D)J and gene expression data. This pipeline generates single-cell feature counts, V(D)J sequences, and annotations for a single library. The mouse reference genome mm10-2020-A was used for aligning gene expression sequences, and the mouse reference GRCm38 v5.0.0 was used for aligning V(D)J sequences. These output files were further used for downstream analysis. This analysis was performed in Harvard Medical School’s high-performance research computing cluster O2.

### Analysis of single-cell gene expression data

All gene expression analyses were performed using R version 4.1.0 [52] and Seurat version 4.1.0 [53]. Other R packages used during this analysis include dplyr version 1.0.8, patchwork version 1.1.1, data.table version 1.14.2, ggplot2 version 3.3.5, cowplot version 1.1.1, viridis version 0.6.2, gridExtra version 2.3, RColorBrewer version 1.1.2, and tibble version 3.1.6. Seurat object was created individually for all samples using the barcode.tsv, features.tsv and matrix.mtx files generated from the *cellranger multi* pipeline. The min.cells parameter was set to 3 and the min.features parameter was set to 200 during the creation of Seurat objects. Cells were filtered out from the Seurat object based on several quality control parameters. First, low-quality cells were removed based on *CD3* gene expression, overall mitochondrial gene expression, and an aberrant high count of genes. We kept the cells if they expressed (>0) either *CD3E*, *CD3D* or *CD3G*, with feature count 200-5000 and overall mitochondrial gene expression less than 20%. Second, we filtered cells based on the expression of housekeeping genes. The list of mouse housekeeping genes was obtained from Satija laboratory website and we kepts cells if they had expressed (>0) at least half of the housekeeping genes in the list. Finally, cells were filtered based on the expression of a list of mouse mitochondrial genes [54]. We removed cells if the mitochondrial gene expression was higher than 2 standard deviations from the mean.

The quality-filtered gene expression data were normalized and scaled by Seurat function *NormalizeData* and *ScaleData* with default “LogNormalize” parameters, generating log-transformed gene expression measurement per 10,000 reads. We next selected the top 2000 highly variable features by implementing the *FindVariableFeatures* function. Using the variable features as input, we performed linear dimensionality reduction analysis using the RunPCA function with 50 principal components (PCs). To create an ‘integrated’ data assay from multiple samples for downstream analysis, we used SCTransformation. The features were selected by *SelectIntegrationFeatures* function for integration. First, using a list of Seurat objects as input, we identified anchors using the *FindIntegrationAnchors* function. Then we integrated the data using the *IntegrateData* function. The integrated Seurat objects were further used for downstream analysis.

### T cell clonal analysis

Matching clones of T cells between blood and islets were determined based on the similarity of TCR sequences. We only included the T cells with at least one alpha and one beta chain in the clonal analysis. The cells were assigned to a particular clonotype if they shared the same amino acid sequence of TCR alpha and TCR beta sequence. This same definition was also applied to cells that carry multiple alpha and beta sequences. Cells having identical TCR alpha or beta sequences were defined as matching clones between blood and islets. For example, if a T cell in blood carries identical TCR alpha or TCR beta sequence to a T cell in islets, it is defined as islet-matching T cell in blood or blood-matching T cell in islets. The total count of cells in a particular clonotype was used to determine the clonal expansion. TCR clonotype data were added to the GEX Seurat object as metadata for integrated gene expression and TCR analysis.

### Cell clustering and psuedotime analysis

The Seurat [53] package was used to calculate UMAP clusters from PCA data. Weighted PCAs were calculated with weights applied to relevant immune markers taken from multiple sources [55–57]. Clusters were determined from PC 1-10 using a resolution of 1.5 and annotated manually by expression of previously determined immune markers (Figure 1e). Monocle3 [58] was used to calculate psuedotime across clusters.

### Weighted co-expression analysis

Genes were clustered by similar patterns of expression across cells using the WCGNA package [59]. Co-expression clusters were generated using a soft power of 10 and a minimum module size of 10, with a merge cut height of 0.15. Clusters were randomly assigned colors to label modules.

### Gene set enrichment analysis (GSEA)

For GSEA, we downloaded the list of Hallmark pathways (v7.5.1) from the Molecular Signature Database (MSigDB, http://software.broadinstitute.org/gsea/msigdb/index.jsp), and used the ranked list of differentially expressed genes between different cell types. R package fgsea version 1.20.0 was used for GSEA.

### Machine learning for the prediction of the matching status of cells

We first extracted the metadata, raw counts, and log normalized raw counts of gene expression from the integrated dataset to predict the matching status of cells based on gene expression profile. Then, we used logistic regression to make predictions with a liblinear solver and lasso penalty of 11. We used the Scikit-learn package in Python version 3.10.4 [60] for the regression analysis and used matplotlib to generate the plots.

### Statistical analysis

All statistical analyses were performed with R or GraphPad Prism software, and P values <0.05 were considered statistically significant. Figure legends show respective statistical analyses.

## List of Supplementary Materials

**Supplementary Figure 1.** Quality control of 5’ gene expression (GEX) library.

**Supplementary Figure 2.** Quality control of VDJ library.

**Supplementary Figure 3.** Feature plots showing the expression of marker genes across T cell clusters.

**Supplementary Figure 4.** An animation shows TCR clonal expansion in the different T cell populations.

**Supplementary Figure 5.** Weighted gene co-expression network analysis (WCGNA).

**Supplementary Figure 6.** Expression of IL-17a by T cells in NOD mice. a

**Supplementary file 1.** Sample metadata for the individual mouse.

**Supplementary file 2.** CD3+ T cell recovery and sequence yield from 10X single-cell library.

**Supplementary File 3.** Expression of marker genes across T cell clusters

**Supplementary File 4.** Differential expression of genes between the islets-matching cells in the blood and blood-matching cells in the islets.

**Supplementary File 5.** Differential expression of genes between islets-matching and non-matching cells in the blood. Positive Log fold-change shows enriched in matching.

**Supplementary File 6.** Differential expression of genes between blood-matching and non-matching cells in islets.

## Acknowledgments

We acknowledge Jennifer Hollister-Lock and the Joslin Islet Isolation Core for helping the purification of mouse islets. We are thankful to Qiong Zhou at Joslin Molecular Phenotyping and Genotyping Core and Stephen Wood at 10X Genomics for providing support during single-cell library preparation. We acknowledge Joslin animal physiology core for the support during NOD mice rearing. MZI acknowledges support for postdoctoral fellowships from the Lundbeck Foundation, Copenhagen, Denmark.

## Author contributions

Conceptualization: ADK, JML, MZI; Mouse experiment, PBMC and islets isolation: MZI; T cell isolation and single-cell library preparation: MZI; Computational and statistical analysis: MZI, SZ, MR, JML, ADK; First manuscript draft: MZI; Manuscript revision: MZI, SZ, MR, JML, JOM, ADK.

## Data and materials availability

All required data are incorporated in the manuscript, supplementary materials, and GitHub repository. Additional information or data requests should be directed to the corresponding author. The single-cell gene expression data were submitted to the Gene Expression Omnibus (GEO) database with an accession number of **GSE200695**. Bioinformatic pipelines and code used in this study are available on GitHub at https://github.com/szimmerman92/NOD_t1d_islets_pbmc_single_cell_tcr

