## Supplementary figures and images for "Single-cell RNA-seq reveals TCR clonal expansion and a high frequency of transcriptionally distinct double-negative T cells in NOD mice"

### Supplementary Figure 1

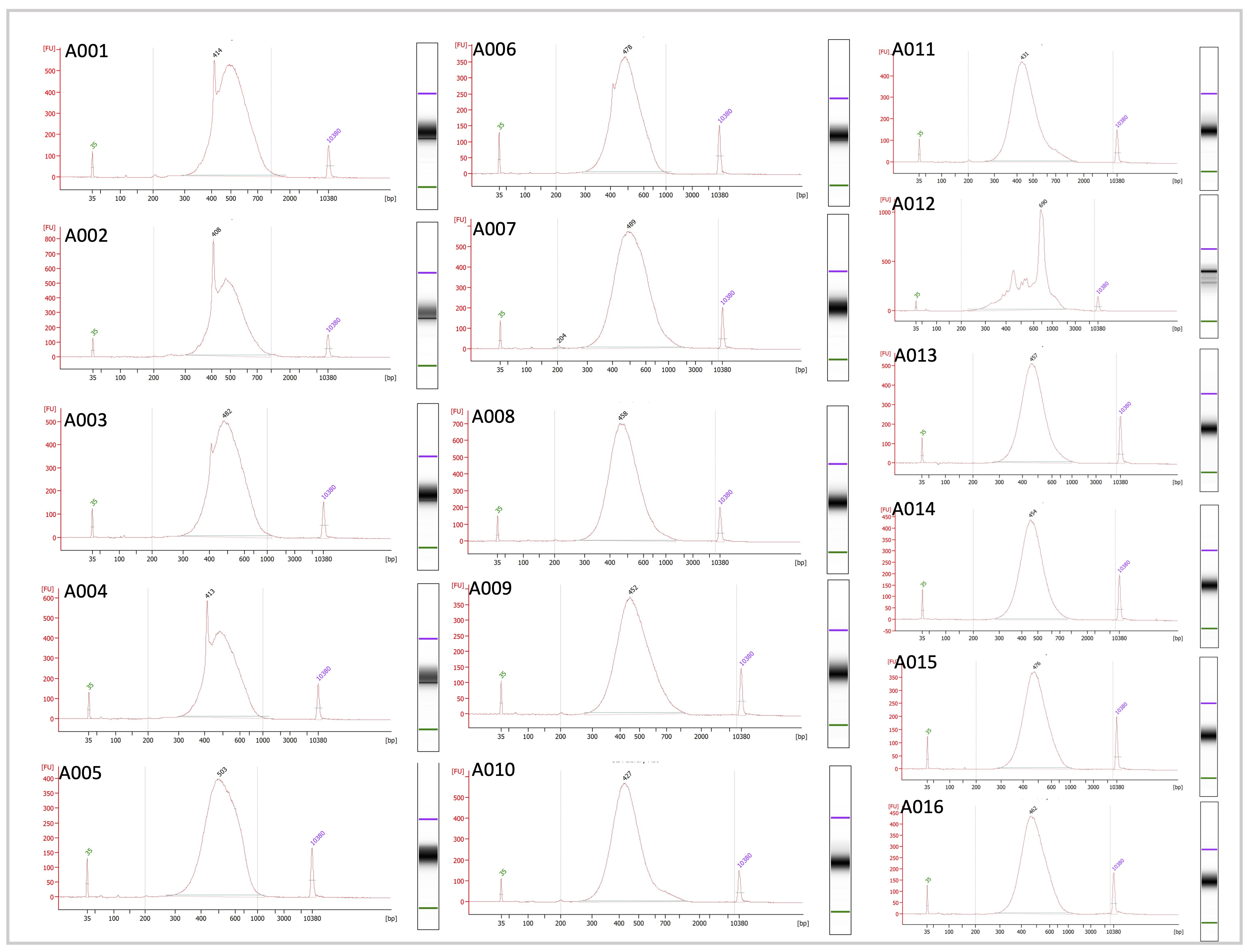

### Supplementary Figure 2

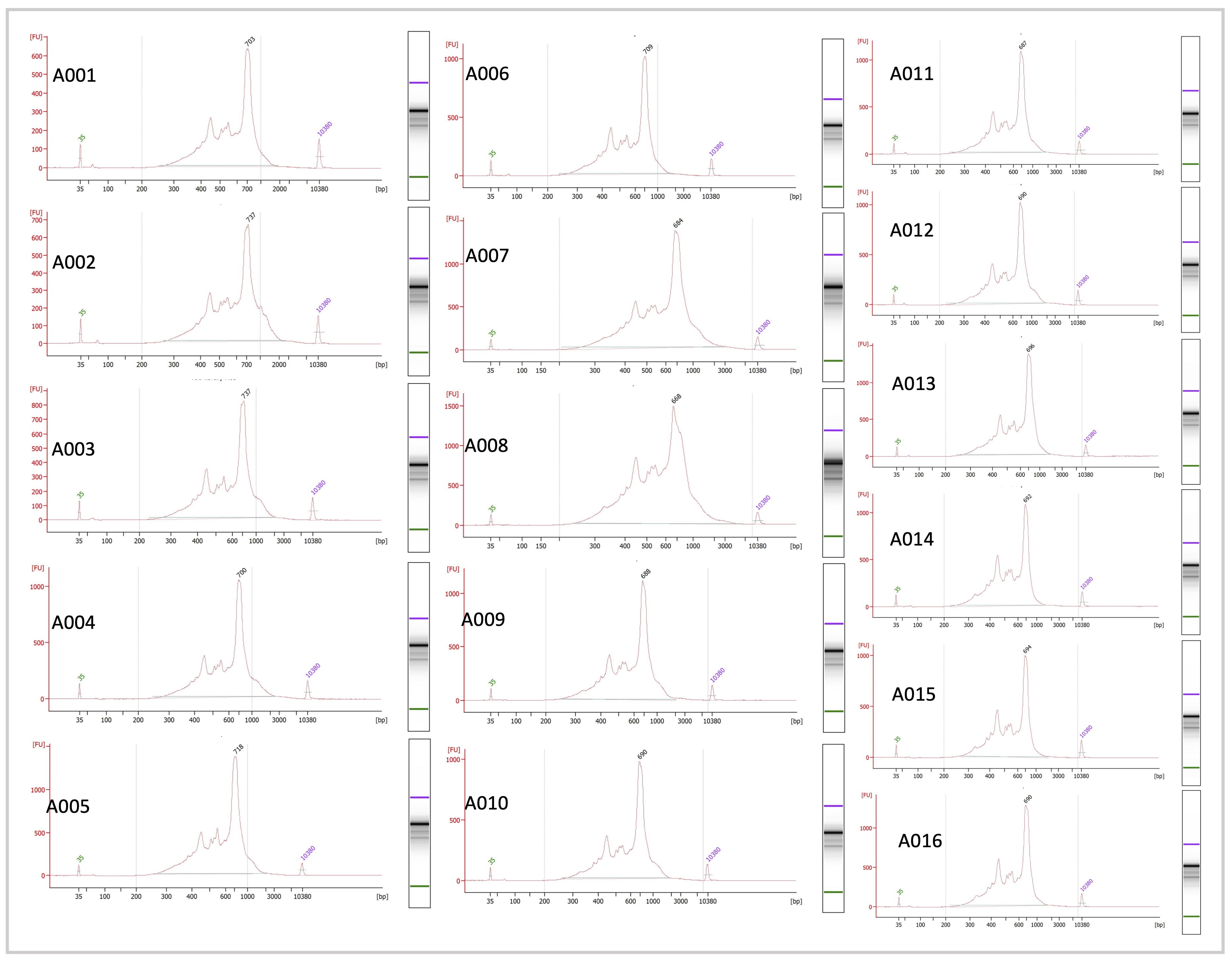

### Supplementary Figure 3

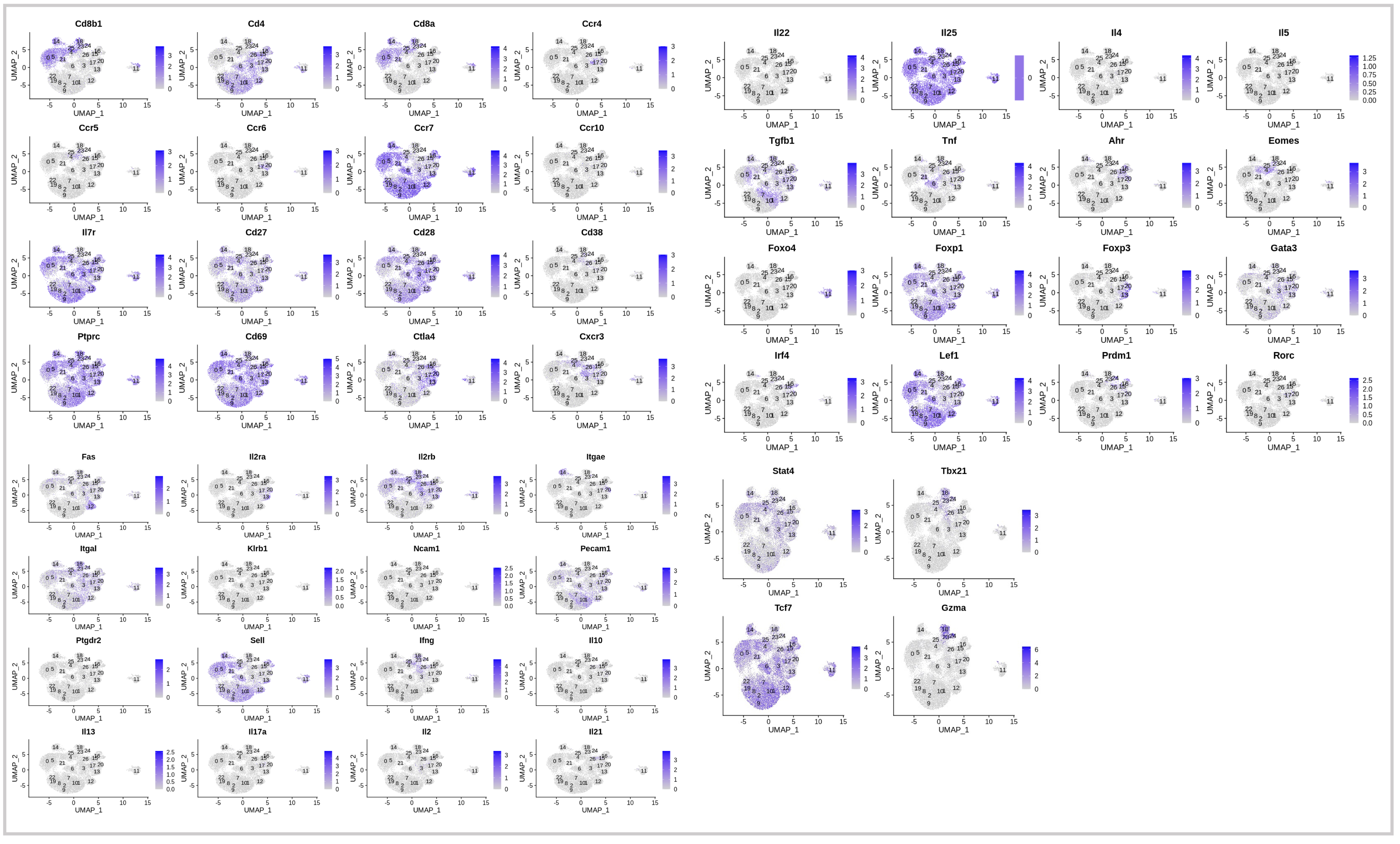

### Supplementary Figure 4

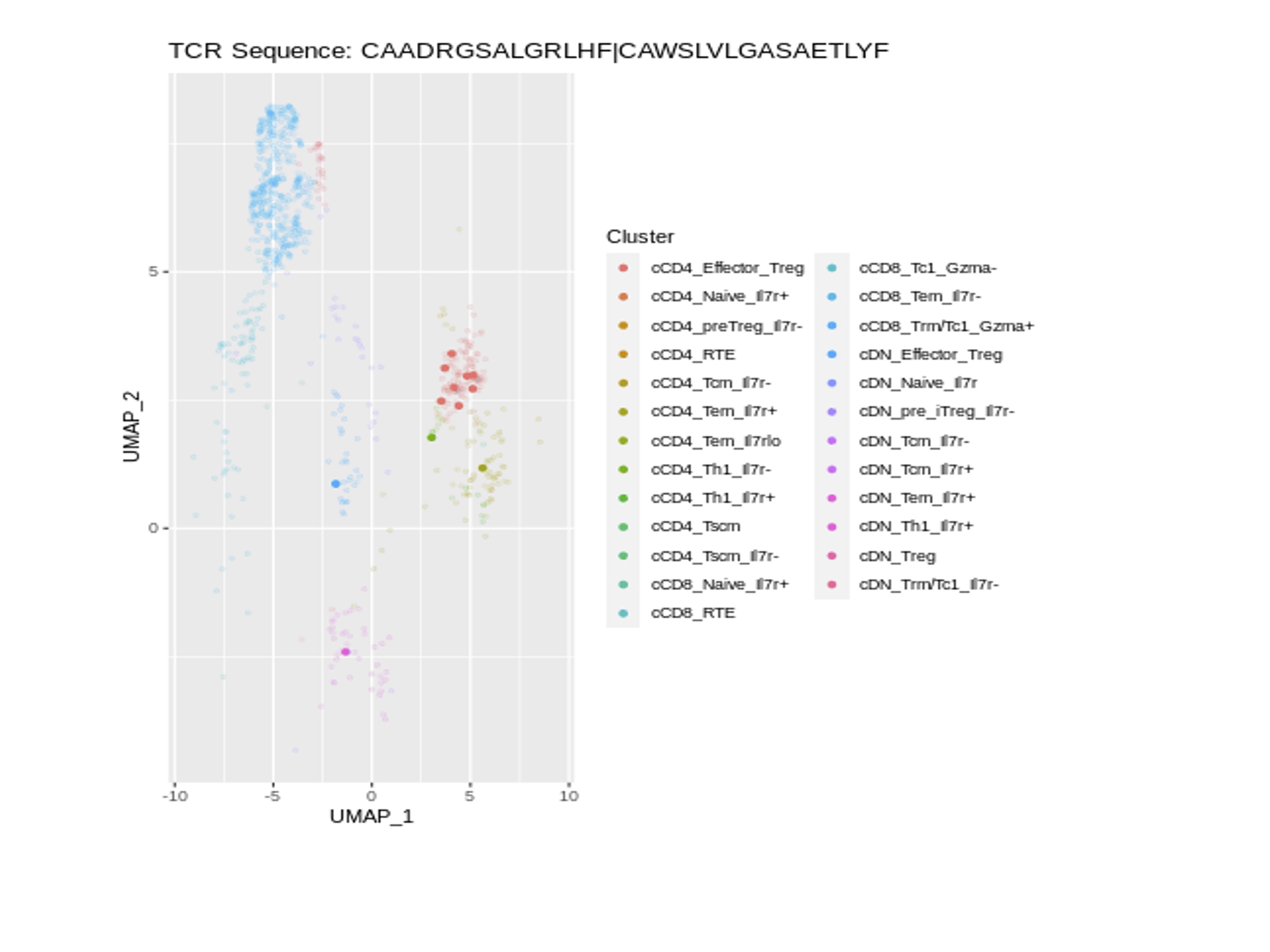

### Supplementary Figure 5

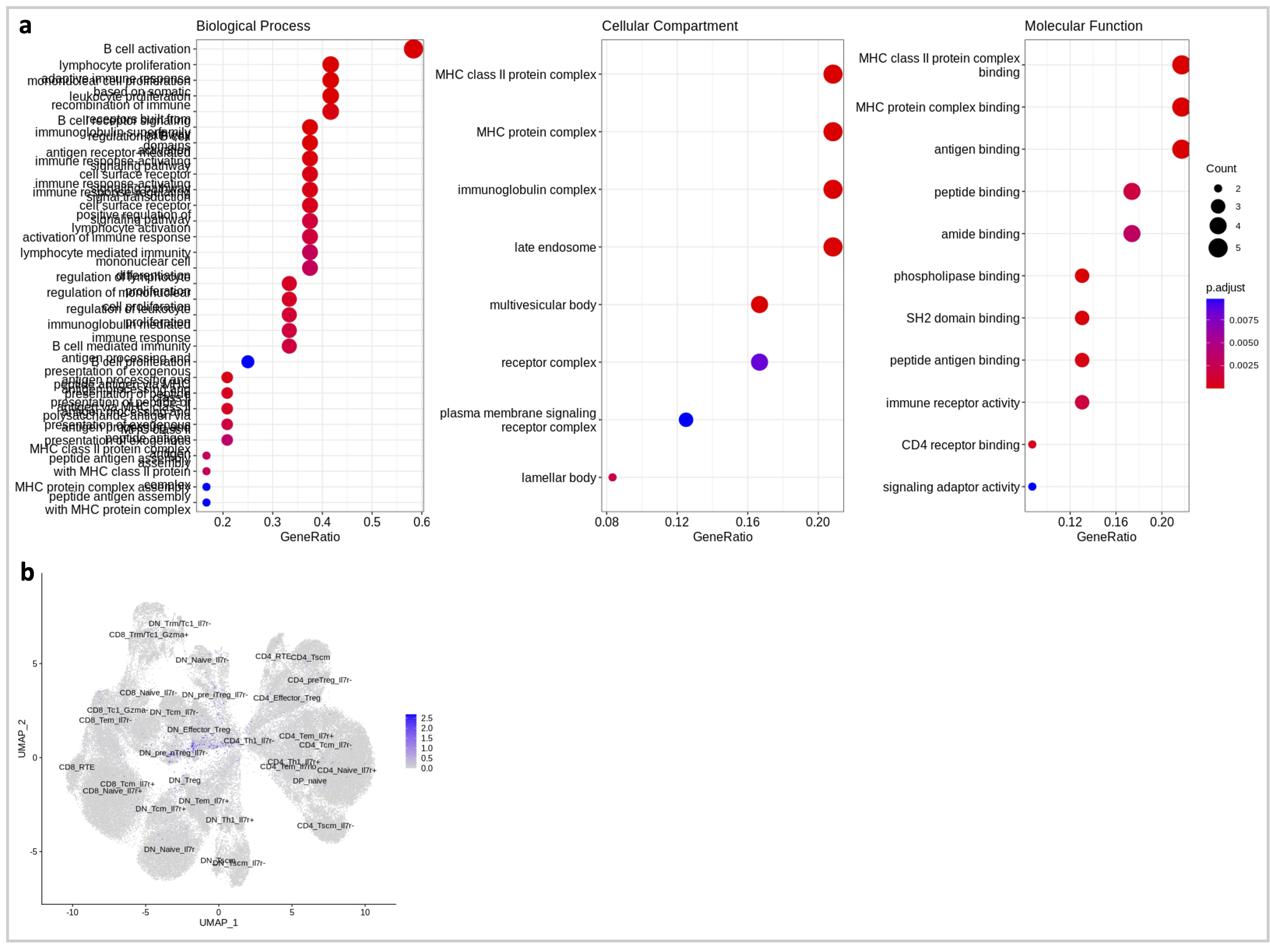

### Supplementary Figure 6

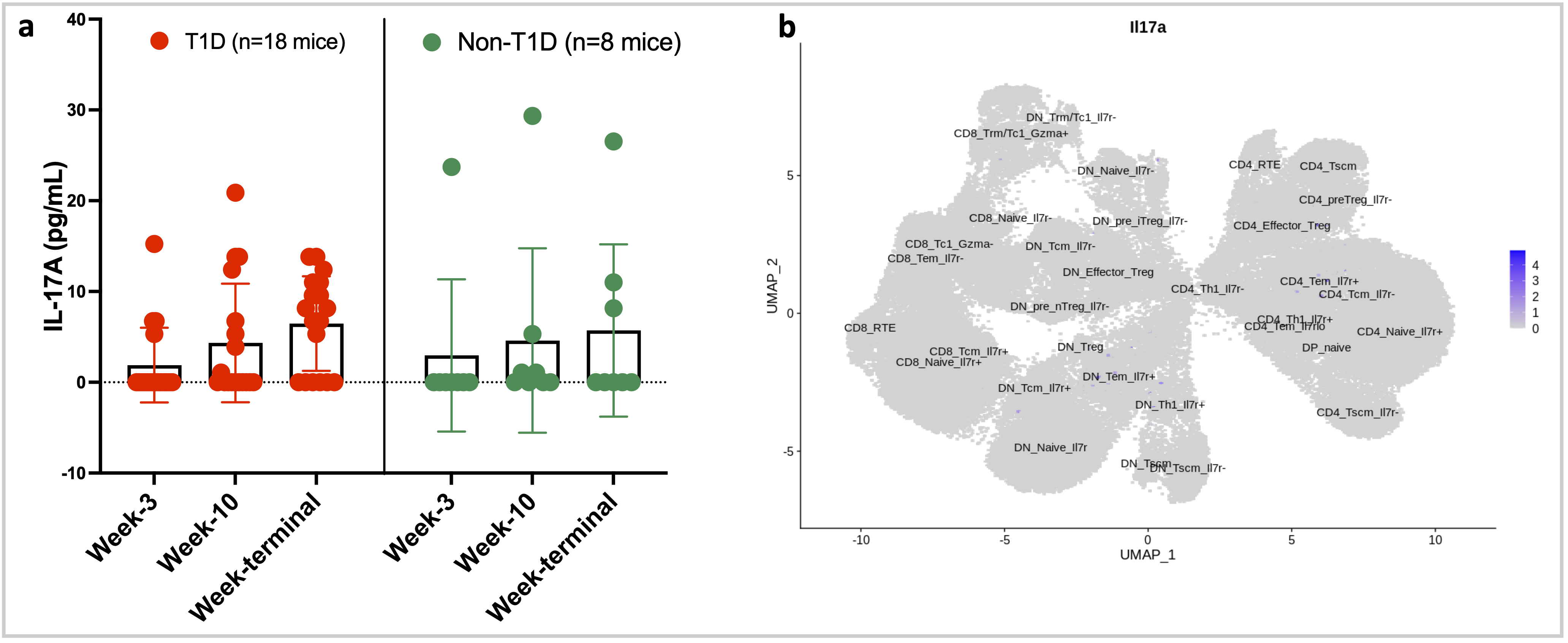
